# Additive effects of environmental and demographic variation shape the repeatability of evolution across replicated experiments

**DOI:** 10.1101/2025.11.03.686244

**Authors:** Karen Bisschop, Meike T. Wortel, Ken Kraaijeveld, Janine Mariën, Isabel Rathmann, Dries Bonte, Steven Declerck, Marjon de Vos, Rampal S. Etienne, Steven Goossens, Jan Kammenga, Frederik Mortier, Joost Riksen, Maurijn van der Zee, Koen Verhoeven, Suzanne Wiezer, Lars Zandbergen, Martijn Egas, Jacintha Ellers, Astrid T. Groot, Marcel E. Visser, Thomas Blankers

## Abstract

The repeatability of evolution is fundamentally important for understanding the origin and diversification of life as well as for developing evolutionary forecasting tools. Repeatability is limited by stochasticity, here defined as changes that are independent of genotypic fitness effects. Over short timescales, the two main sources of stochasticity of evolutionary change are environmental stochasticity and demographic (life-history) stochasticity. Quantifying the relative importance of these two sources of stochasticity in driving fitness outcomes is crucially important for predicting evolutionary responses. To gain insights in the effects of stochasticity, five institutes replicated an evolutionary experiment exposing *Caenorhabditis elegans* to novel rearing conditions. Replication across the institutes led to variation in selective environments, e.g. through divergent microbiomes among institutes. Replication within institutes was done across demographic treatments that influence the potential for population-size dependent fluctuations in allele frequencies (drift) and genetic hitchhiking (draft). We found high among-institute variation in fitness outcomes, which was partially explained by variation in microbiota. Whereas lab-specific effects explained most of the variance in mean fitness, the repeatability of fitness outcomes depended more on demographic heterogeneity. Specifically, population bottlenecks resulted in high among-replicate variation in fitness. When combined, environmental and demographic stochasticity additively reduced repeatability, underlining their additive importance in developing evolutionary forecasting tools. These results further highlight the importance of statistically integrating heterogeneity in experimental evolution to identify factors constraining outcome repeatability and study replicability.

## Introduction

The predictability of evolution has attracted major interest in evolutionary biology [1–3]. Quantifying the extent to which evolutionary theory can be predictive and identifying the factors that contribute to predictability help us to better understand the origin and present-day patterns of biodiversity, such as whether convergent phenotypes (i.e., similar traits in divergent lineages) result from convergent natural selection or from evolutionary constraints and chance events [4,5]. At the same time, identifying the mechanisms influencing evolutionary predictability will result in more accurate and more broadly applicable forecasting [3]. However, accurately predicting evolutionary trajectories and outcomes poses substantial challenges, even when predicting short-term evolutionary responses, such as forecasting trait change or fitness outcomes in response to the rapid, contemporary changes in our climate.

One major challenge is to accurately estimate the strength of selection. Selection strength depends on a complex interplay of variables, including resource and trait distributions, genetic architectures and species interactions [3,6]. Overcoming this challenge is possible by collecting more accurate data on organisms and their environment. A second major challenge is to account for evolutionary historical contingencies, such as historical population structure, that have shaped the genomic background of the targets of selection [7,8]. Both challenges are characterised by stochasticity, i.e. changes in genomes, individuals, and environments independent of adaptation [9]. Stochasticity inherently limits the predictability of evolution, for example by random changes in the available genetic variants or in the fitness of these variants. Evolutionary genetic changes may always be highly sensitive to stochasticity, resulting in low repeatability of genetic trajectories [1]. Evolutionary forecasting of fitness should be more accurate, but it is unclear how sensitive predictions are to stochasticity and contingencies in evolution. Here, we aim to measure the sensitivity of end-point repeatability of fitness (which we here refer to as the fitness response) to stochasticity. Although we recognise that repeatability is not same as predictability, quantifying the effect of stochasticity on the repeatability of fitness evolution will uncover inherent limitations to developing functional predictive models and help identify possible solutions.

Three major forms of stochasticity in evolution are mutational stochasticity, environmental stochasticity, and demographic (life history) stochasticity [9]. When predicting short-term evolutionary change that mostly depends on standing genetic variation, mutational stochasticity can be largely ignored. Environmental stochasticity results from spatio-temporal changes in interactions with the abiotic and biotic environment and results in variation in the selective regime. Studies on adaptive evolution in wild populations have demonstrated that spatio-temporal fluctuations in selective pressures are a major source of long-term unpredictability [10,11]. In the lab, random fluctuations in selection through time have been found to impact microbial evolution by diversifying the direction and magnitude of the selection response [12–14].

Demographic (or life history) stochasticity affects the variation available to selection and the genetic background of selected variants and results from individual-level stochastic events. These events, such as survival and reproduction, are described by a probability that is scaled by population size (drift). Similarly, recombination and the segregation of genotypes are described by probabilities that depend on linkage and genetic hitchhiking (draft) [9]. Evolutionary experiments have shown that bottlenecks or other factors that influence the population size can limit repeatability because drift increasingly drives evolution when effective population is low [15–20] leading to divergent evolutionary trajectories over time (Lenski and Travisano 1994). However, as long as population sizes are not excessively small (effective population size, Ne > 10) and selection is not excessively weak (selection coefficient, s > 0.01), variation in selection will be more important than drift for trait repeatability and predictability [22,23]. In contrast, draft is important at normal/high population sizes, in particular when recombination is low and in the presence of sign epistasis [24–26]. Sign epistasis is when the direction (sign) of the fitness effect of a variant depends on other variants in the genome and is commonly observed in nature [26].

Concordantly, much of the variance in trait evolution and fitness outcomes can be attributed to variation in founder populations [8,27] and differences in repeatability across experiments are strongly driven by patterns of recombination and linkage in the genome [27–29]. These findings show that environmental differences and historical contingencies can strongly influence evolutionary outcomes. Both factors arise from demographic heterogeneity across experimental and natural populations and affect the repeatability of trait and fitness evolution. Whereas environmental stochasticity influences selective regimes, demographic stochasticity influences selection responses. Disentangling the relative importance of these sources of stochasticity for the repeatability of evolution will inform what data to collect and how to analyse this data to increase the accuracy of evolutionary predictions.

### Rationale of the experiment

In this study, we measure the relative importance of environmental and demographic stochasticity in shaping repeatability of fitness outcomes of micro-evolution (i.e., adaptation within species from standing genetic variation). Experimental evolution provides a potent tool to determine the causes of repeated evolution as well as to investigate under which circumstances repeatability is lacking [30]. In our evolutionary experiment, we introduced a multiparent intercrossed *C. elegans* population [31,32], to which we refer as the ancestor, to a novel environment. The changes relative to the ancestral environment include: 1) a temperature change, from 20° C to 16° C; and 2) a novel food source, i.e., the bacterium *Priestia megaterium* instead of *Escherichia coli*. The experiment was replicated in five different institutes, with detailed outlined instructions. Despite these instructions, small differences originated across the participating institutes, leading to undirected environmental heterogeneity. These variations arose from factors such as different handling routines during nematode transfers and treatments, as well as divergent microbiota shaped by lab-specific environmental microbiomes that became associated with the nematodes over time (see Results). Thus, the temperature-change, feeding on *P. megaterium*, and institute-specific conditions jointly made up the selective environment, which varies from institute to institute (Fig. 1) and creates environmental heterogeneity. We also introduced secondary treatments that modulated demographic processes and, consequentially, the potential for drift and draft. These treatments included variation in starting population size (500, 50, or 5; population bottleneck), reproductive mode (obligate outbreeding, facultative self-fertilisation, or obligate self-fertilisation), population age synchronisation through bleaching, and reduced gene flow via isolation. The evolutionary experiment spanned 15 weeks, corresponding to approximately 20 generations. This duration is generally sufficient to detect measurable changes in fitness, as demonstrated in previous experimental evolution studies [33].

**Figure 1.**
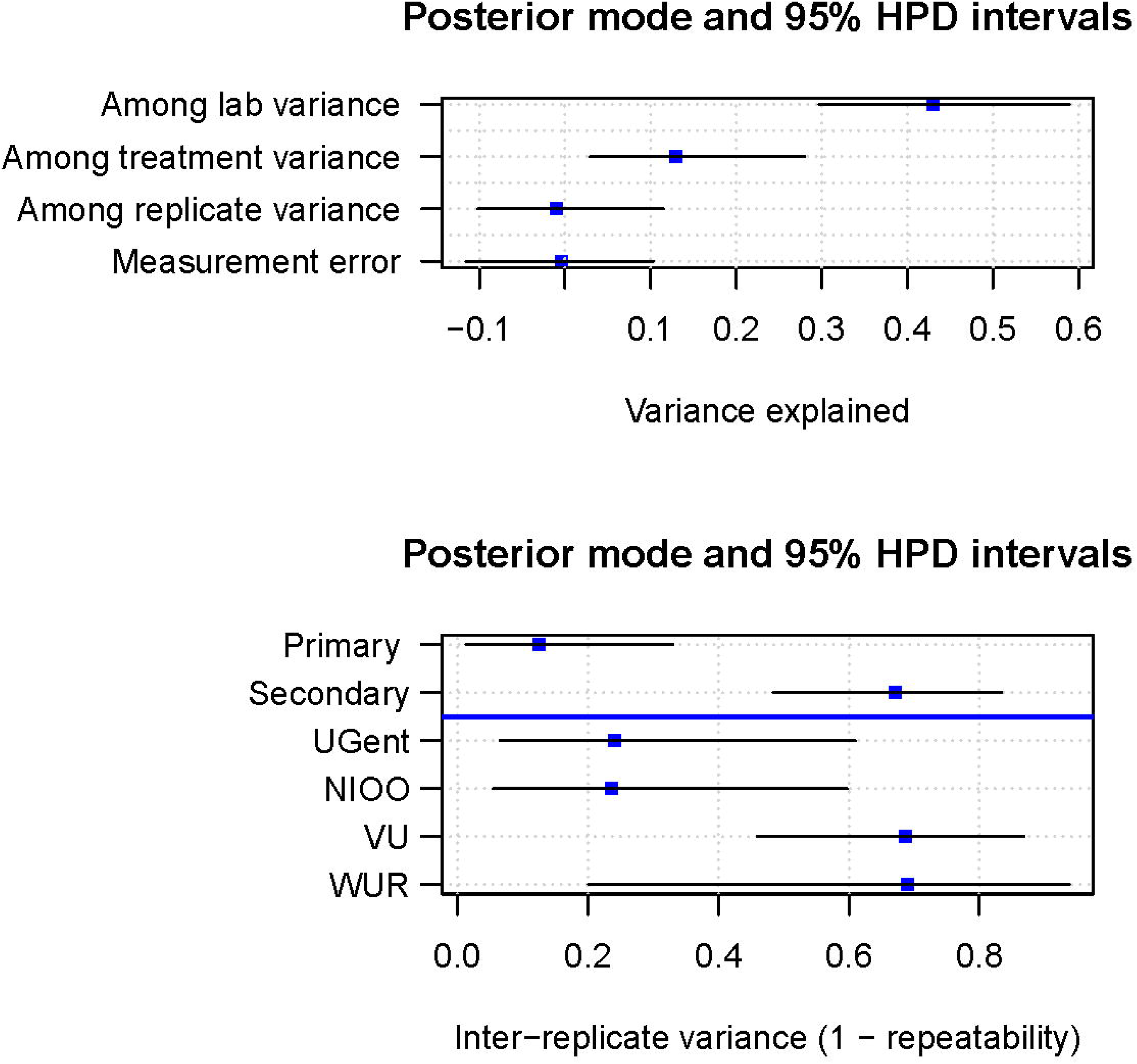
Environmental and demographic stochasticity. All institutes (institute A and B) performed an identical evolutionary experiment where *C. elegans* and microbial communities evolved on a novel resource, the primary treatment (Tr. 1). The controlled conditions during the experiments (temperature and other climate attributes, as well as food source) were kept similar (indicated with the thermometer symbol), but the uncontrolled conditions were different (e.g., institute microbiota in grey and blue, and institute handling routines as symbolised with the hand icons), resulting in environmental heterogeneity across institutes. Furthermore, all institutes except one conducted a second treatment (Tr. 2) which differed between institutes and resulted in demographic heterogeneity across treatments.

We first compared fitness in *C. elegans* before and after the 15-week evolution across experimental conditions and research institutes to quantify sensitivity of evolutionary outcomes to environmental and demographic heterogeneity. We approximated fitness by measuring *C. elegans* population growth rate on *P. megaterium* at 16°C. Population growth rate is driven both by developmental rate and fertility and increasing growth rates over time are thus interpreted as adaptation to the novel conditions. Measuring growth over 7 days corresponds to a single rearing step in the evolutionary experiment. Subsequently, we sequenced the nematodes and associated microbiota using metagenomic sequencing. In this study, where we focus on evolutionary responses manifesting in fitness changes, we leverage these metagenomic data to assess environmental variation (reflected in the microbiota) and to examine if variation in population growth rate at the end of experimental evolution depends on microbial diversity. Lastly, we modelled the heterogeneity of the evolutionary experiment in Bayesian mixed effect models to decompose the effect of environmental stochasticity (across-institute variability) and demographic stochasticity (within-institute variability) in influencing the repeatability of the response in population growth rates. We expected highest repeatability within institutes within treatment, and progressively lower repeatability when comparing across treatments, institutes, or both. In line with this expectation, we expected institute-specific microbiota to influence variation in population growth rates.

### Repeatability and replicability of experimental evolution

In addition to *repeatability* of evolution, this study also informs on the *replicability* and external validity of evolutionary experiments in the context of environmental and demographic stochasticity. We define *repeatability* as the degree to which independent evolutionary processes lead to similar genetic, phenotypic, or fitness outcomes when subjected to similar selective environments. *Replicability* is defined as the degree to which independent researchers, using the same methods but different datasets, arrive at the same conclusions (National Academies of Sciences (2019).

Importantly, a lack of replicability in the fields of animal behaviour, ecology, and experimental evolution, has recently put emphasis on systematic introduction of heterogeneity in genetic and environmental factors [35–38] to develop more robust theories in ecology and evolution that do justice to the complexity of biological systems. In evolutionary experiments that explore the repeatability of evolutionary processes, historical contingency can cause divergent outcomes at the level of genotype, phenotype, or fitness [8]. Such studies thus particularly benefit from including *a priori* conceived [38] or *a posteriori* characterised heterogeneity [39] at the level of organisms and their environment. Ultimately, incorporating heterogeneity across institutes uncovers how reliable evolutionary inferences in one institute translate to a broader context (i.e., the external validity) [38].

## Results

### Adaptive evolution in the primary treatment

We first investigated whether the final (week 15) populations of *C. elegans* increased in population growth rate compared to the starting populations (week 1) at the five research institutes (Figs. 2A and 2B). Overall, variance in population growth rate was explained by institute (*χ*² = 127.2 and *P* value < 0.0001) and the interaction between institute and week (*χ*² = 32.0 and *P* value < 0.0001). Two institutes evolved higher population growth rates (UG: *z* ratio = 3.6 and *P* value = 0.0003; VU: *z* ratio = 2.4 and *P* value = 0.0186) and one institute lower growth rates (UGent: *z* ratio = -3.6 and *P* value = 0.0003) across all replicates from week 1 to week 15 (Fig. 2; Table S2).

**Figure 2.**
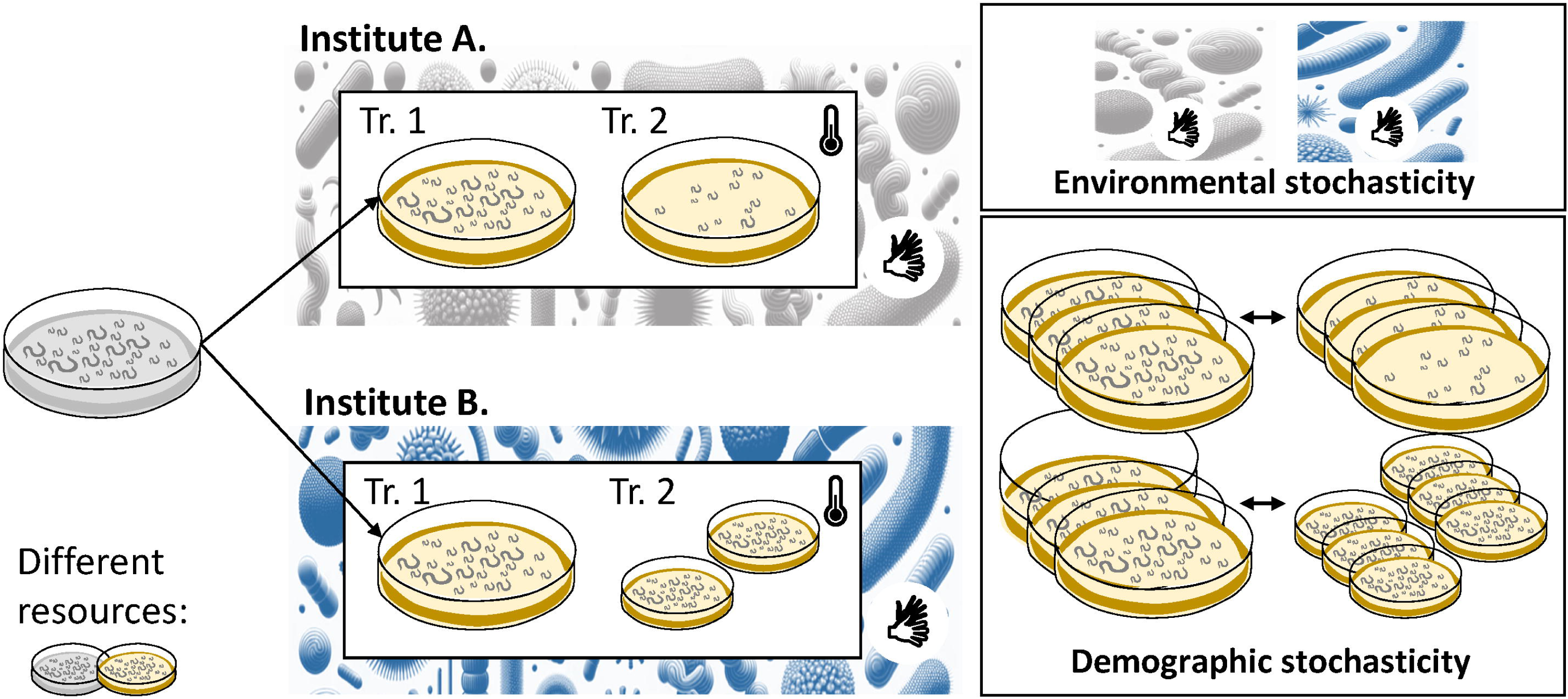
Growth rate evolution. **A)** Daily growth is calculated as 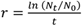 with t = 7 and N0 = 500. Solid points are per-biological replicate average growth rates (Anc.: n = 5, NIOO: n = 5, but data for one replicate is missing in week 1; UG n = 5, UGent n = 5; VU n = 5; WUR n = 3) across the technical replicates (ranging from 3 to 9 per biological replicate). Lines connect week 1 and week 15 averages for the same replicate. Boxes show inter-quantile range, median; whiskers are 1.5 x interquartile range of the spread in daily growth across the technical replicates (the grey solid point is an outlier from the boxplot). **B)** Selection response per replicate line is calculated as r_week15_ – r_week1_. Box-and-whiskers are as in A). **C-F)** The y-axis shows the daily growth rates, and the colours indicate the time point during experimental evolution (white: week 0, light grey: week 1, and dark grey: week 15). Solid points are per-biological replicate average growth rates and lines connect biological replicates across time points when multiple time points are available. The effect of population bottlenecks with the different treatment shown in different panels: no bottleneck, moderate bottleneck (50 individuals at the start), or strong bottleneck (5 individuals at the start) is shown in C. The effect of bleaching with the left panel without initial bleaching step and the right panel with initial bleaching step is shown in D. The effect of reproductive mode with the three secondary treatments shown as different panels: dioecious, androdioecious, and monoecious is shown in E. The effect of mixing with the mixing of three petri dishes during each refreshment step or the isolation of the petri dishes is shown in E.

### Adaptive evolution in the secondary treatment

Population growth rates varied among institutes (Figs. 2A-B) and among treatments (Figs. 2C-F). The final growth rate was higher in the monoecious populations, in the isolated and in the bleached treatments compared to dioecious (*t* ratio = -2.8 and *P* value = 0.0218), mixed (*t* ratio = -8.4 and *P* value < 0.0001), and not-bleached (*t* ratio = -4.9 and *P* value = 0.0004) populations in the primary treatment (Figs. 2C-F, Table S4).

### Environmental stochasticity – institute-specific microbiomes

By replicating the experiment across different institutes, we introduced undirected environmental heterogeneity that was independent of the *C. elegans* genotype. At least two sources of variation contributed to this heterogeneity. The first were small, undirected fluctuations in the protocols used in transfer and treatment of the nematodes (Appendix I). The second were undirected differences in the microbial communities that became associated with the nematodes over time.

To assess the microbial community composition and diversity, we leveraged the metagenome sequence data obtained for 93 ancestral, before, and after evolution lines. Prior to sequencing, repeated washing of the sampled nematodes removed most environmental bacteria and fungi. Microbial sequences obtained here are thus assumed to be from bacteria and fungi intimately associated with the nematodes. Between 1% and 91% of trimmed reads were classified as bacterial or fungal genomes (Appendix II). Microbiomes identified in sequence pools through metagenomic profiling showed an increase in alpha diversity, meaning higher species richness, over time for three institutes: UGent, UG, and VU (Fig. S3A, Table S3) and were divergent among institutes (Figs. S2 and S3B).

### Marginal effects of microbiome variation on population growth rates

We tested whether the random variation in environment affected population growth rate by modelling the daily growth rate as a response of institute, treatment, and microbiome composition (using its principal components). We identified week, institute, and microbiome PC3 (but not microbiome PC1 and PC2) as predictors for growth rate variation in the set of samples subject to the primary treatment only (ANOVA: *F*_1_ = 5.35; *P* value = 0.026; when adding PC3 to linear model, ΔAIC = 3.92; Table S5). The bacterial genera mainly driving PC3 are *Serratia*, *Pseudomonas*, and *Brucella* and a total of 8.72% of the variance in microbiome composition across samples is explained by PC3. Using the same data, we tested whether microbiome variation explained variation in population growth rate *change* (from week 1 to week 15) using a random forest regression. Presence/absence of specific taxa (relative abundance >1% in at least one sample) predicted growth rate change variation in addition to institute specific effects (Fig. S4). In the set of samples that also included all secondary treatments, including microbiome PC axes improved model fit and when included in the model, PC axes significantly explained population growth rate variation (PC1: *F*_1_ = 5.20; *P* value = 0.026; PC3: *F*_1_ = 5.56; *P* value = 0.021; Table S6).

### Environmental and demographic stochasticity additively and similarly influence repeatability

We investigated the effect of environmental stochasticity (inter-institute variability) versus demographic stochasticity (intra-institute, between-treatment variability) on the (endpoint) repeatability of population growth rates. For this we fitted generalised linear mixed effect models in an MCMC framework using the R-package MCMCglmm [40]. We first sequentially added terms (institute, treatment, replicate ID, measurement) to a model explaining daily population growth variation. Only treatment was fitted as a fixed effect because that was a statistically non-random variable. Most of the population growth rate variation (∼50%) was explained by differences among the institutes, followed by differences among treatments (∼10%; Fig. 3A). We then used these models to estimate the relative among-replicate variance (which is 1 – Repeatability) across institutes within the primary treatment only, across institutes within all treatments, and across treatments within each institute. Repeatability was constrained more by demographic variation than by environmental variation. Among-replicate variance was higher, with non-overlapping 95% HPD intervals, across treatments within VU and WUR versus across institutes (Fig. 3B). Among-replicate variance across treatments within UGent and NIOO was also higher than across institutes, but in this case 95% HPD intervals overlapped (Fig. 3B). Environmental and demographic heterogeneity appeared to mostly additively affect repeatability: inter-replicate variance across secondary treatments and institutes (the line marked secondary in Fig. 3B with posterior mode = 0.68) was similar to the inter-replicate variance across institutes within the primary treatment (posterior mode = 0.10) plus the average of the inter-replicate variances across treatments in each of the institutes (posterior mode = 0.53).

**Figure 3.**
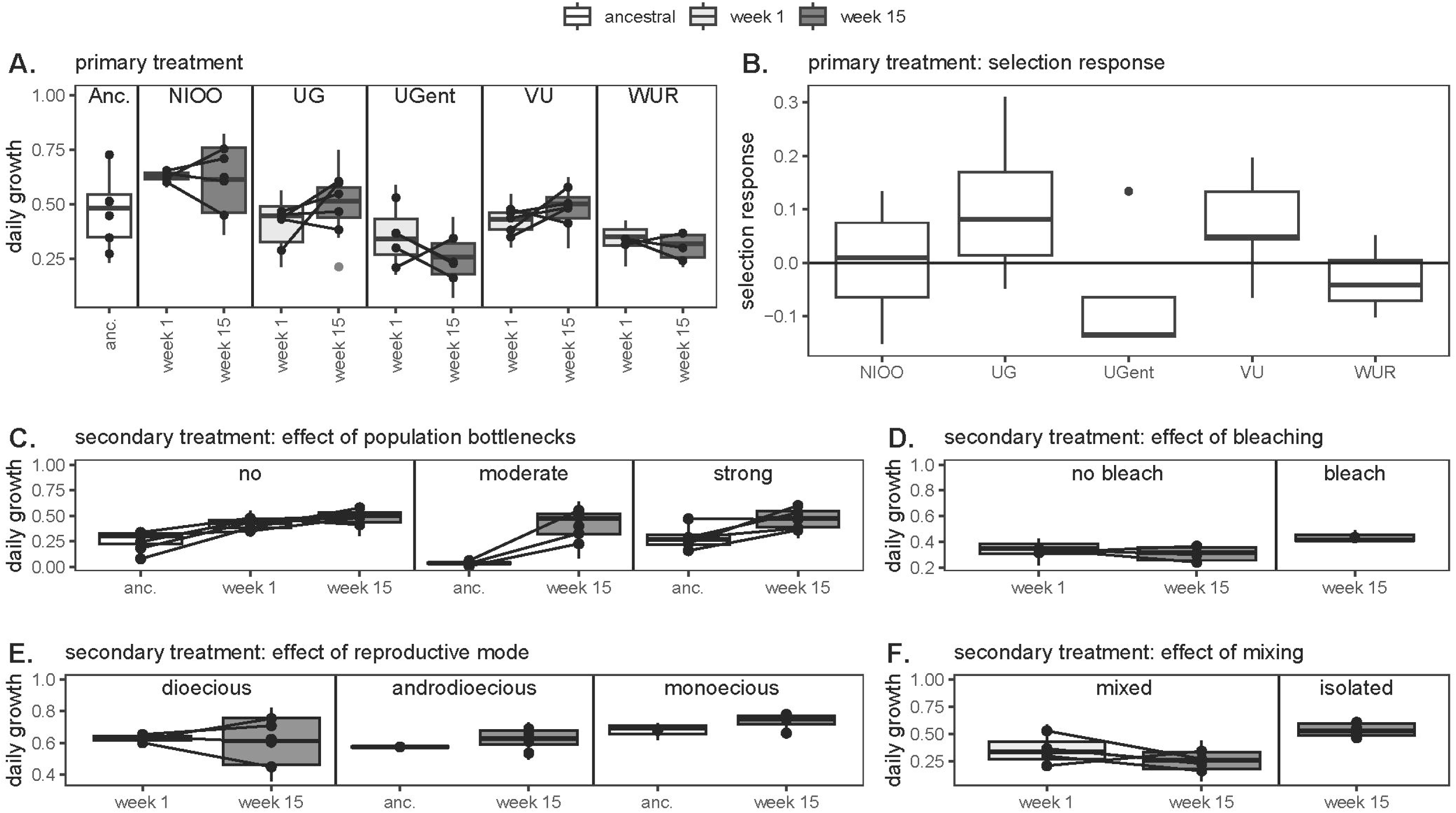
**A)** Population growth rate variance partitioning. Population growth rate variance explained by among-institute variation (environmental stochasticity), among-treatment variation (demographic stochasticity), among-replicate, and among-measurement variation. Blue rectangles indicate the posterior mode, and lines delineate the 95% HPD interval of the posterior distribution for variance components in a MCMC generalised linear model. Some variance estimates are slightly negative, which is mathematically implausible but can occur due to the MCMC model’s partitioning approach and prior specification. This suggests that the estimated contribution to trait differences is negligible. **B)** Population growth rate repeatability. Repeatability (among-replicate variance relative to total variance) of week 15 population growth rate across institutes and across treatments within institutes (UG is not shown as no secondary treatments were performed in this institute). Blue rectangles indicate the posterior mode, and lines delineate the 95% HPD interval of the posterior distribution for variance components in a MCMC generalised linear model.

## Discussion

Parallel evolving populations in nature have shown that both spatiotemporal heterogeneity in the environment and heterogeneity in evolutionary and demographic history limit the repeatability of evolution [4,5,10,11]. Without experimental work that quantifies the effect of demographic and environmental heterogeneity, it remains unclear to what extent evolution is repeatable across ecological contexts [3] and to what extent evolutionary predictions can be optimised by more accurate measurements of selection and by accounting for population genetic structure [2,6,23]. We thus conducted an evolutionary experiment across multiple institutes in The Netherlands and Belgium that introduced heterogeneity in the environment and demography of evolving populations. This approach allowed us to quantify random limits on the repeatability of evolution and measure the replicability of evolutionary experiments across heterogeneous contexts.

Nematodes in only two out of five institutes consistently evolved higher population growth rates on *P. megaterium* and under the lower temperature relative to their ancestor (Fig. 2). Most of the variation in population growth rates is attributable to institute differences (Fig. 4A). Environmental heterogeneity across institutes likely resulted in different selective regimes, as evidenced by the different communities of microbiota (Fig S3) with a weak but non-zero effect on population growth rate evolution (Fig S4, Table S5,6). Within institutes, population genetic and demographic properties were varied. The treatments likely caused an inflation of life-history variation relative to the primary treatment alone. This variation in reproductive modes, bottlenecks, demographic distribution, and the random mixing of local genotypes influences several aspects of evolutionary change, including the probability that a given genotype transfers to the next generation (i.e., drift), the rate of inbreeding (i.e., local drift), and the strength of linkage between selected and neutral loci (i.e., draft) [41]. Together, environmental and demographic treatments introduced heterogeneity that we quantified as among-replicate variation in our mixed model framework. We conclude that both environmental and demographic heterogeneity affect repeatability and that they do so additively. Demographic stochasticity appears to have stronger effects on repeatability than environmental stochasticity, depending on the specifics of the treatment.

This means that incorporating more detailed information on variation in selection and tuning to the life history specifics of focal populations subject to predictions is important for forecasting evolution. Moreover, stochasticity inherent to evolutionary processes acting across heterogeneous environments and populations inevitably introduces noise, resulting in substantial individual variation and often low explanatory power, even when predicting seemingly redundant traits such as fitness over brief time scales.

### Variation in fitness means among institutes and treatments

The temporal change in growth rates across institutes, increasing in two institutes, two staying roughly the same, and one decreasing, reflects multiple possible processes. Identifying divergent microbiota among institutes confirmed one aspect of the expected heterogeneity in selection environments. Because the central experimental paradigm involved adaptation to a novel bacterial food source, we expected the microbial background environment to be relevant for the adaptive process. Even though our analyses revealed some statistically significant effects from microbiome diversity on fitness and fitness change through time, we conclude that these effects are either weak or institute specific. An alternative, non-mutually exclusive explanation for the among-institute variation is that adaptation to the experimental conditions in one institute may translate poorly to fitness measured in a different institute. Growth rates for the evolved populations from all institutes were measured at VU. This institute was one of the sites where population growth rates increased over time. Conducting all measurements at a single location ensured that fitness means and variances were independent of the experimenter and equipment. However, it is not unusual that replicate populations with little fitness variation under the conditions in which they evolved exhibit higher fitness variation in other environments [8]. For example, local adaptation and environmental trade-offs may lower the growth rate measured in populations at the VU that have evolved elsewhere [42,43].

The variation in growth rates across treatments depended on multiple factors. As expected, final growth rates were higher in monoecious populations than in dioecious populations, possibly due to a greater proportion of reproductive individuals. Growth rates were also higher in the isolation and bleach treatments compared to the mixed and non-bleached populations. Isolating rather than mixing the three Petri dishes per each biological replicate may be beneficial by reducing outbreeding depression [44] and by reducing migration load [45,46]. By synchronising developmental stages and minimising founder effects, the bleach treatment may have enhanced the expression and selection of alleles beneficial early in life, thereby maintaining adaptive potential. Compared to institute differences, which we assume affected selection coefficients, treatments likely mostly affected population genetic structure. Overall, we found institute differences to explain most of the variation in population growth rates, which is likely due to a combination of the divergent background microbiota and lab-specific adaptive processes. Treatments explained less variation in mean growth rates, after accounting for institute differences.

### Environmental and demographic heterogeneity additively impact fitness repeatability

The D00 ancestor of our experimental populations harboured high levels of genetic variation, showed a rapid decay of linkage disequilibrium (LD) over short genetic distances, and exhibited a high degree of sign epistasis, all as a consequence of the multi-parental hybridisation [31,32,47]. Selection likely acts on many different loci that may be expected to be involved in feeding on specific microbial prey. In a context of high genetic redundancy and extensive epistasis, the extent to which populations may ‘find’ the same fitness-improving solutions within the limited time of approximately 20 generations likely depends on several factors. It is shaped not only by selection but especially by the genetic background and genetic structure within which selection operates [26,48]. In line with this expectation, we show an important role for demographic stochasticity. This finding highlights that predictability of adaptive evolution across different loci is inherently limited by the contingencies associated with founder populations, echoing previous results from studies in microbes, flies, and nematodes [8,27]. A forthcoming analysis of the genetic selection response can reveal whether variation in LD and epistasis across the genome in response to the demographic treatments may indeed explain the observed effects on repeatability of fitness outcomes. The current results on the repeatability of fitness outcomes imply that both sources of stochasticity must be considered when predicting short-term adaptive responses to environmental change. This stochasticity likely introduces substantial noise, reducing the accuracy of such predictions.

### Replicability and heterogeneity in experimental evolution

An important advantage of our approach is that we do not limit our analysis of repeatability to a single institute or a universally shared demographic and genetic background. Most experimental evolution studies are conducted in a single institute, using standardised media and equipment. These studies typically apply either constant selection coefficients (i.e., constant selection) or constant selection differentials, where the coefficient increases proportionally with the selection response [49]. Although this ensures precise estimates of repeatability, it does not reflect the broader biological reality of evolution occurring across different ecological and demographic contexts. Whenever spatiotemporal heterogeneity is introduced, evolutionary outcomes diverge among replicates that differ in environment more than among replicates that experience similar conditions. This is evident from our results and supported by findings from both highly controlled and more ‘natural’ experiments. For example, seemingly minor differences in media, glassware, and atmospheric conditions can substantially influence the outcomes of microbial evolutionary experiments [14,38,50]. Only 3 out of 80 quantitative trait loci for biomass traits in rapeseed (*Brassica napus L.*) were stable across two experimental locations [51], and just 53 out of 152 behavioural responses in mice were reproducible across five institutes [52]. Moreover, the population of origin (genetic background) can itself be a major driving force, as shown both in this study and in previous experimental evolution work using diverse founders [29,53,54]. Our study illustrates how environmental and demographic heterogeneity can be incorporated to increase the external validity, which makes experimental results more generalisable beyond a single laboratory setting.

In our data, we assessed among-replicate variance across similar conditions (within-institute, within-treatment), across intermediate dissimilar conditions (replicates across institutes *or* treatments), and across most dissimilar conditions (replicates across institute *and* treatments). We find that repeatability decreases accordingly and is not uniformly dependent on whether replicates vary by treatment or by institute. Statistically integrating heterogeneity in experimental evolution through the mixed model approach helped identify factors and quantify their influence on evolutionary repeatability and experimental replicability. We thus were able to compare different sources of heterogeneity quantitively to consider how they impact the uncertainty associated with extrapolating findings to broader contexts. Testing repeatability across heterogeneous contexts aids in understanding (the limits to) evolutionary predictability. Replicating evolutionary experiments across locations and experimenters, and including heterogeneous contexts, adds broader biological context and probes the external validity of replays of the tape of life.

## Materials and Methods

### Study species and resource

The bacterivorous soil nematode *Caenorhabditis elegans* (Maupas, 1900) is highly suitable for experimental evolution given its small body size (adult females are approximately 1 mm), high fecundity (about 300 eggs per adult female per week), short generation time (can be as short as 50h), and easiness to maintain in the laboratory [55]. Furthermore, frozen records can be created given that the first larval stage can undergo cryopreservation [55]. The *C. elegans* D00 population [31,32] served as the initial source population for the entire experiment. The D00 population is a dioecious, multiparent, intercrossed population that harbours high levels of genetic variation maintained by obligate outcrossing [31,32] between males and females in a 1:1 sex ratio.

The D00 population was shared by the Teotónio laboratory (IBENS-Paris) and expanded on *Escherichia coli* OP50 at 20°C at the University of Amsterdam (UvA) to divide and distribute the starting populations in aliquots (hereafter referred to as ‘ancestral’ population). Each institute received ten frozen aliquots of this starting population, which then evolved on a novel food bacterium, the gram-positive bacterium *P. megaterium* (DSM No. 509 – formerly known as *Bacillus megaterium*). Both *E. coli* OP50 and *P. megaterium* were distributed from the same starting population and maintained at 4°C in the different institutes. Bacterial lawns were added on the nematode growth medium (NGM) plates by pipetting 50 µL of bacteria in L broth and spreading it with a sterile hook or sterile loop. Rearing, media preparation, washing nematodes off plates, and freezing protocols mostly followed standards [56] but protocols are available in Appendix I.

### Primary treatment

#### Shared axis of selection

In the primary treatments, obligate outcrossing *C. elegans* populations were allowed to evolve on *P*. *megaterium* for 15 weeks, at 16°C (Fig. S1). All populations of *C. elegans* were maintained on ±12 mL NGM agar plates in Petri dishes (94 x 16 mm, sterile with vents, Greiner Bio-One International GmbH). These NGM plates were poured at the Netherlands Institute of Ecology [NIOO] using a Petri dish filling machine (Mediajet and Mediaclave by INTEGRA Biosciences Ltd. Thatcham, Berks RG19 4EP, UK), distributed among the five participating institutes (NIOO, University of Groningen [UG], Ghent University [UGent], Vrije Universiteit Amsterdam [VU], and Wageningen University [WUR]), and stored in the institutes afterwards. We noticed from 16S rRNA amplicon sequence data coming from buffer washes over the plates that all these plates were contaminated with a (non-visibly growing) *Stenotrophomonas sp.,* which potentially indicated an additional but equal selection pressure on the evolving populations. The other shared axes of selection thus were: 1) The resource change (i.e., *P. megaterium* instead of the standard *E. coli* OP50 diet) was the primary focus; the new food source, *P. megaterium*, is known to be hard-to-eat, delays growth, and is non-toxic [50 and personal observations]. And 2) the temperature change (i.e., 16°C instead of 20°C) was chosen to obtain about one nematode generation per week; in this way all experimental procedures could be performed on weekdays. However, the effects of the two perturbations and plate contamination cannot be disentangled and are together considered as the shared (across institutes) axis of selection.

At the start of the experiment, in each institute ten aliquots of the ancestral population (Fig. S1) were defrosted by holding the cryo-tubes in the hand. When thawed, the liquid was directly pipetted onto an NGM plate. All ten aliquots were combined as starting population except at WUR where only six aliquots were used and at VU where each replicate started from a single aliquot. This initial step was performed with *E. coli* OP50 as a food source to avoid selection during the expansion. Afterwards the nematodes were collected by washing the different plates and the density of nematodes was estimated. The evolutionary experiment was initiated on fresh NGM plates with a lawn of *P. megaterium* under equal numbers of nematodes per replicate per institute (in two institutes, UGent and UG, the initial population size was 500 nematodes, whereas it was 1000 for the other institutes). For each replicate (i.e., five replicates per institute except at WUR where only three replicates were done) three plates were maintained per replicate to sustain more genetic variation, support larger population sizes, and to avoid the loss of a replicate if one plate would fail. To ensure these benefits, 500 nematodes were transferred by washing the plates, mixing the three plates per replicate, estimating the nematode density, and pipetting the necessary volume for transferring 500 nematodes to the new plates. The remaining nematodes were cryopreserved for use in population growth rate assessment and DNA extraction.

#### Environmental stochasticity – variation in handling routines

In addition to the novel conditions shared among institutes, there was environmental heterogeneity across institutes. That variation manifested in the following main variables: (i) differences in institute handling routines and (ii) institute-specific microbiota. The differences in institute handling routines can be grouped in differences in institute set-up (e.g. the use of a flow cabinet or flame, and the presence of light in the climate cabinet or climate room), the preparation of the plates with bacterial lawns (e.g. the location where NGM plates without bacteria were stored, the time the bacteria could grow on the plates, and the rate at which new bacterial cultures were prepared), and the washing of the nematodes off the plates (e.g. the use of M9 or S buffer, the volume of buffer used, and the time the buffer was on the plate). All identified differences are listed in Appendix I.

#### Environmental stochasticity - microbiome profiling

To profile institute-specific microbiota associated with C. elegans, we aimed to sequence all replicates from primary and secondary treatments at both the start (week 0 or 1) and end (week 15) of the experiment. However, due to sample limitations, sequencing was not successful at week 0/1 for WUR secondary and UGent secondary treatments, and for one replicate each in UGent and UG primary treatments. DNA was extracted from several hundreds of individuals per population using the DNeasy B&T kit (Qiagen) after washing the sample three times with M9. The population pools of DNA were sequenced by IGAtech (Udine, Italy) on the Illumina Hi-Seq platform using 150 bp paired-end reads. Quality of the reads was checked with fastQC and poor-quality nucleotides at the ends of the reads were trimmed using trimmomatic. Quality-checked and trimmed reads were mapped against the WBCel235 reference genome [58,59] using the Burrows-Wheeler Aligner [60]. Reads that mapped against the reference genome were filtered out using samtools. The remaining reads in the alignment were converted back to fastq files. We performed taxonomic classification of the non-host sequencing reads at the genus level with the k-mer based kraken2 software [61] and obtained species abundances with Bracken [62]. The database (PlusPf database, latest version 9/4/2024) included bacterial, viral, archaea, protozoa, and fungi references. Genus levels that did not reach 1% abundance in any of the samples were filtered out, leaving 31 different genera in total. For further analysis, species abundances were Hellinger transformed to remove unit-sum constraints of compositional data and principal component analysis as well as redundancy analysis was performed with the rda function of the vegan package 2.6-10 in R [63] to reduce the number of dimensions and analyse composite multi-species dimensions in relation to institute and treatment variation. The loadings in the PCA were generated with the vegan::scores function with scaling two.

### Secondary treatments

#### Demographic stochasticity

In the secondary treatments, four different variables were considered: the influence of a population bottleneck, the effect of mixing within-population replicates, the influence of different reproductive modes, and the effect of an initial bleaching step (Fig. S1). Each parameter was varied at only one specific institute, and all other conditions were as in the primary treatment.

At VU, population bottlenecks were introduced in the starting populations prior to the first week of experimental evolution. Therefore, only the initial population size in week zero differed from the primary treatment. Three treatment levels were investigated: a strong bottleneck (population created from five founders), a moderate bottleneck (population created from 50 founders), or no bottleneck (population created from 500 founders). The expectation was that smaller effective population sizes would increase the potential for drift.

In the default protocol, the three different Petri dishes per replicate are mixed prior to transfer (Fig. S1), which constitutes a form of migration among sub-populations. In the isolation treatment at UGent five Petri dishes were kept isolated instead resulting in five replicates. In this isolation treatment, overall levels of genetic variation may be reduced, and locally adaptive alleles (local within a single Petri dish) are not recombined among plates.

At NIOO, two alternative reproduction modes were contrasted with the obligate dioecious D00 that was used in the primary treatment: obligately self-fertilising populations (monoecious populations or M00) and wildtype populations (androdioecious or A00; Theologidis *et al.*, 2014). In the wildtype populations, sexual reproduction occurred through self-fertilisation and through the production of males that cross with hermaphrodites. The populations are all derived from the same multi-parent hybrid ancestor, A00, into which loss-of-female-spermatogenesis [fog-2[q71)] and male-killing [xol-1(mt3055)] alleles were introgressed to create the D00 and M00 populations, respectively [31,47]. We expect the monoecious population to exhibit a higher growth rate than the dioecious population, because twice as many individuals can reproduce. Also, we expect that the obligate dioecious population will maintain the highest genetic diversity and has the highest adaptive potential. In contrast, the monoecious population may adapt rapidly but face evolutionary constraints due to reduced genetic variation. The performance of the androdioecious population is expected to fall between that of obligate outcrossers and selfers, depending on its outcrossing rate. If male frequency increases under early stress and declines as conditions improve, A00 may initially resemble D00 but shift toward wild-type sex ratios over time. This plasticity could allow A00 to benefit from both mating systems. However, if male frequencies cannot respond quickly to selection, the population may instead experience disadvantages of both systems (e.g., reduced mating opportunities or inefficient resource allocation). Thus, A00’s outcome might depend on the extent and timing of its outcrossing response.

At WUR, an initial bleaching step was done prior to the onset of the experimental evolution. Bleaching is a common procedure in *C. elegans* rearing and kills all organisms (including contaminant bacteria) except the unhatched embryos inside the gravid females. Consequently, it is used to sterilise, and age synchronise populations. It is expected that this treatment mostly affects the demography in the population (as all individuals have the same age at the start of the experiment). Synchronising the population at the embryonic stage may favour alleles beneficial early in development while reducing founder effects, as reproduction is shared across the entire cohort. This could improve adaptive potential by maintaining broader genetic representation. An overview of the treatments, treatment levels and sample sizes can be found in Table S1.

### Population growth rate assessments

Ancestral populations were cryopreserved after expansion in the lab, but before exposure to the novel conditions. Starting populations were cryopreserved after one week of growth on *P. megaterium* at 16°C. Final populations were cryopreserved after 15 transfers/weeks of growth on *P. megaterium*. The population growth rate of all populations was assessed at a single location (VU Amsterdam) to prevent assessment bias. Cryopreserved samples were thawed and reared on *E. coli* at 20°C for one week (∼two generations) to create a common garden, remove (grand)parental effects that could prime the offspring for the new environment, and obtain sufficient nematodes for the start of the assessment. This assessment thus compares heritable, (epi)genetic effects on population growth rates across replicates and conditions. Population growth rate was assessed for three technical replicates, each starting with 500 nematodes, per replicate unless the expansion step did not yield enough nematodes. For some replicates, the assessment was repeated on different days (resulting in more technical replicates: 3 x the number of measuring days). The population size was estimated by washing nematodes off the plate using 2 mL M9, collecting the nematode suspension in a microcentrifuge tube and extrapolating from the number of nematodes counted in three droplets of 5 µL well-mixed nematode suspension. The counts after seven days are used to calculate the growth rate, as this was the only timepoint available for all populations. The growth rate was calculated by taking the logarithm of the population size at day 7 divided by the population size at day 0 (500) (i.e., the initial population size at day 0), which was then divided by seven to have a daily growth rate 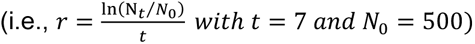.

Growth curves obtained for populations that also had counts at earlier timepoints (4, 5, 6 days) were used to validate growth rate calculations at this timepoint (see Appendix I). Our manual counts were highly correlated with counts from a BioSorter flow cytometer following linear regression (Appendix I). Some samples at UGent showed strong visual contamination, which reduced the growth of the nematodes. These samples could therefore not be included in the further analyses. We also removed samples with low survival of nematodes after freezing, which was the case for five out of 138 sample measurements.

### Statistical analysis

#### Population growth rates in the primary treatment

Unless otherwise mentioned, statistical analysis was done in R version 4.4.2 [64]. Two samples from UGent that showed strong contamination apparently limiting growth and showed up as outliers in statistical distributions were removed. We evaluated whether daily population growth rate differs between the populations after one week and after 15 weeks in each of the institutes using GLMMs (generalised linear mixed models) with glmmTMB package version 1.1.10 [65] with a Gaussian distribution with identity link (Shapiro-Wilk normality test: *W* = 0.9928, *p*-value = 0.4329). The dependent variable was the growth rate, and the week (categorical variable: week 1 or week 15), the institute (categorical variable: NIOO, UG, UGent, VU, and WUR), and their interaction were modelled as explanatory variables. The random variables included in the model were the date on which population growth rate assessments were conducted and the replicate ID (to link the measurements on the same replicate across weeks). Residual diagnostics were performed using the DHARMa package [66], and no deviations from model assumptions or other issues were detected. We did a Wald χ2 test to assess the statistical support for including fixed effects and performed pairwise comparisons (with emmeans v1.10.6; Lenth, 2024), adjusted for multiple comparisons with Tukey’s method. We calculated the selection response as the difference in population growth rate between week 1 and week 15 populations.

#### Population growth rates in the secondary treatment

We examined whether daily population growth rate varied between secondary treatments within each institute after 15 weeks using generalised linear mixed models (GLMMs) implemented in the glmmTMB package v1.1.10 [65]. A Gaussian distribution was applied for UGent (Shapiro-Wilk normality test: *W* = 0.96662, *p*-value = 0.1681) and WUR (Shapiro-Wilk normality test: *W* = 0.95634, *p*-value = 0.6291). For the VU and NIOO institutes, we used an orderNorm transformation with the bestNormalize package v1.9.1 (Peterson & Cavanaugh, 2020). The model’s dependent variable was growth rate, and the independent variables were the secondary treatments. Date of assessments and replicate ID within secondary treatments were included as random effects. Residual diagnostics were assessed using the DHARMa package [66]. Overall, most models did not show significant deviations from uniformity, dispersion, or outliers. However, for two out of five institutes, minor issues were detected: in one model, some quantile regressions failed, and in another, the Levene test indicated significant heterogeneity of variance. These deviations were considered minor and not expected to substantially affect the interpretation of the results. Pairwise comparisons were conducted using the emmeans package v1.10.6 (Lenth, 2024), with Tukey’s method applied to adjust for multiple comparisons.

#### The effect of microbiome on population growth rate

To assess whether variation in microbiota is the result of institute-specific accumulation of environmental microbes, we tested if the diversity of the nematode-associated microbiota increased from week 1 to week 15 in the primary treatment. We estimated the Shannon diversity across microbiota using the function ‘diversity’ in the vegan package v2.6-10 [63] and performed a Kruskal-Wallis rank sum test per institute with week as explanatory variable.

We examined whether variation in population growth rate was explained by microbiome composition, accounting for week and institute. To do this, we fitted linear models (using lme4 v1.1-35.5, Bates *et al.*, 2015) with the daily growth rate (averaged over the repeated measurements for each replicate ID) as response variable. Explanatory variables included institute, week, and the first three principal components of microbiome variation. We then used backwards model selection based on likelihood ratio-tests to refine the models. We also assessed if change in the population growth rate from week 1 to week 15 was explained by changes in the microbiome composition through a random forest analysis with ntree = 100000 using randomForest v4.7-1.2 [70]. These analyses were restricted to primary treatment data.

#### Effect of environmental versus demographic stochasticity on repeatability

We investigated the effect of environmental (among-institute variability) and demographic heterogeneity (within-institute, between-treatment variability) on the (endpoint) repeatability of adaptive evolution (population growth rates). We first asked whether week 15 daily growth estimates were more variable among institutes or among treatments within institutes using nested linear models (institute:treatment:replicate:repeated measure) implemented in the R-package MCMCglmm [40], using R version 4.3.1. Hence, both primary and secondary treatments were included in this model. If environmental stochasticity is the main factor constraining the outcome after 15 weeks, we expect more variance to be captured by the institute term compared to the treatment or replicate term, which capture treatment and random effects, respectively. We iteratively added nested variables to the model, starting with institute, and then calculated for each model the increase in variance explained relative to a null model with no explanatory variables.

We then asked whether repeatability depended more on environmental stochasticity versus demographic stochasticity. We used linear mixed effect models in MCMCglmm to compare among-replicate variance in population growth rates attributable to institutes with among-replicate variance attributable to treatments. This was done using separate models per institute. Repeatability was considered to be higher when among-replicate variance in growth rates was lower. If environmental stochasticity is the main factor constraining repeatability, we expect that there is more among-replicate variation partitioned across institutes than across the treatments. We first fitted a global model with treatment as fixed effects and the institute, and technical and biological (evolutionary) replicates as random effects using uninformative priors (degree of belief ν = 0.002). Posterior estimates for the variance among measurements and replicates were then used to define informative priors with prior variance equal to the posterior mode and degree of belief equal to the posterior mode divided by the 95% Highest Posterior Density interval (smaller spread relative to the mean will increase degree of belief). The informative priors were used here to incorporate information from the full data set to help model convergence when analysing small subsets of the data. We fitted the following models: 1) only the primary treatment data with institute as (random) explanatory variable; 2) both primary and secondary treatments with institute and treatment as explanatory variables; 3) models for each institute-specific data with treatment as explanatory variable. We estimated posterior distributions across 1,000 MCMC samples (1,000,000 iterations, 100,000 burn-in, 1,000 thinning interval) and calculated the mode and 95% Highest Posterior Density [71].

## Author Contributions

M.E.V, A.T.G., J.E, M.E., M.T.W., and K.K. designed research; K.B, M.T.W., K.K., J.M., S.G., F.M., J.R., S.W. and L.Z. performed research; K.B., T.B., and I.R. analysed data; K.B. and T.B. wrote the paper with input from all authors.

## Data Availability

Data and R code used for analysis and visualisation are publicly available in Zenodo [https://doi.org/10.5281/zenodo.17497646] and all genetic data have been deposited in the NCBI Sequence Read Archive under BioProject accession [PRJNA1252273].

## Acknowledgments

We would like to thank the Teotónio lab (IBENS-Paris) for providing the nematodes, and J. Teapal and the Utrecht University Large Particle Flow Cytometry Facility (UU-LPC) for their help with the BioSorter. We also acknowledge The Predicting Evolution consortium (M. Bosse, T. de Jager, L. de Jeu, A. de Visser, M. Groenen, A. Heerdink, P. Hogeweg, M. Maan, I. R. Pen, I. Smallegange, S. van der Steen, S. van Doorn, B. Voetdijk, and B. Wertheim), and the Origins Center for helpful discussions. This work was funded by The Dutch Research Council National Science Agenda (NWA-ORC 400.17.606/4175), a Flemish Research Foundation fellowship awarded to KB (FWO 12T5622N), IR was supported by NWA.1418.22.013, and LEZ was supported by the Adaptive Life program of the Faculty of Science and Engineering, within GELIFES.

