## Supplementary information for "Additive effects of environmental and demographic variation shape the repeatability of evolution across replicated experiments"

FIGURES

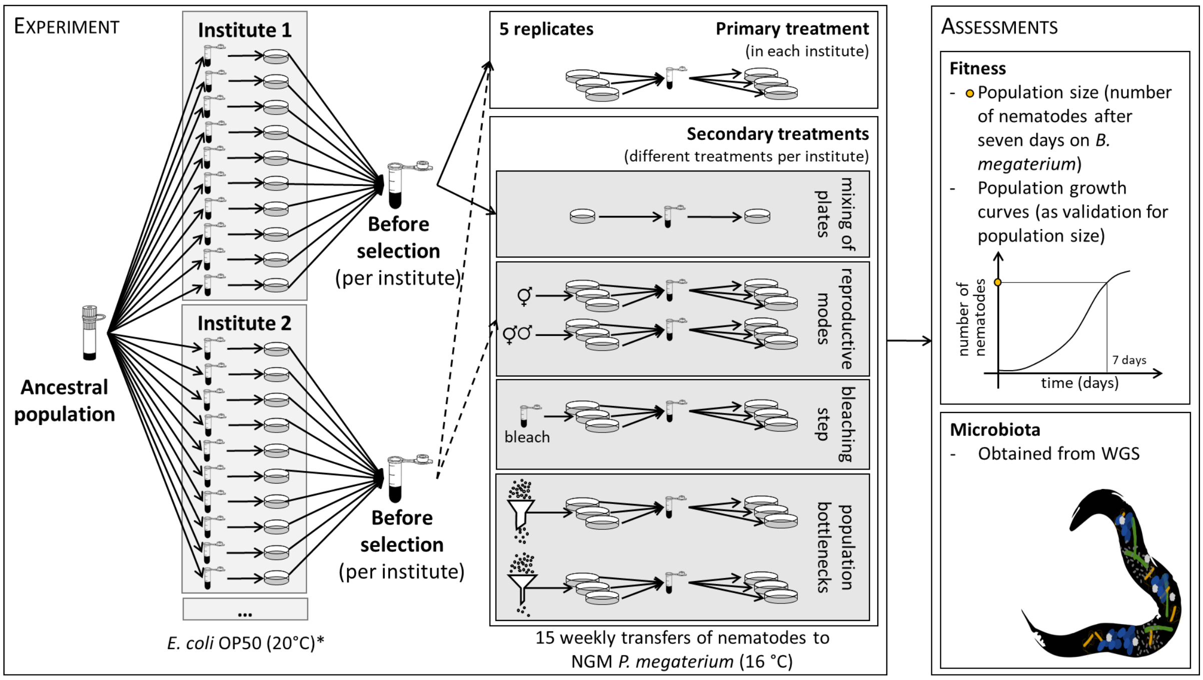


***Figure S1: General overview figure.*** *The left panel is an overview of the experiment itself, where each institute received ten aliquots originating from the same ancestral population (at WUR only six aliquots were used for the before selection population, and at VU each replicate started from a single aliquot). Within each institute a shared experiment (primary treatment) was performed, with fifteen weekly transfers onto fresh NGM* P. megaterium *plates at 16°C. Each institute had multiple replicates, and each replicate consisted of three plates of which the nematodes were mixed every week. Besides, side experiments were performed investigating the influence of secondary treatments: removal of mixing, reproductive modes, an initial bleaching step, or population bottlenecks on the evolutionary experiment. The right panel shows the assessments of fitness and microbiota (*in one institute the expanding temperature was 32°C instead of 20°C).*

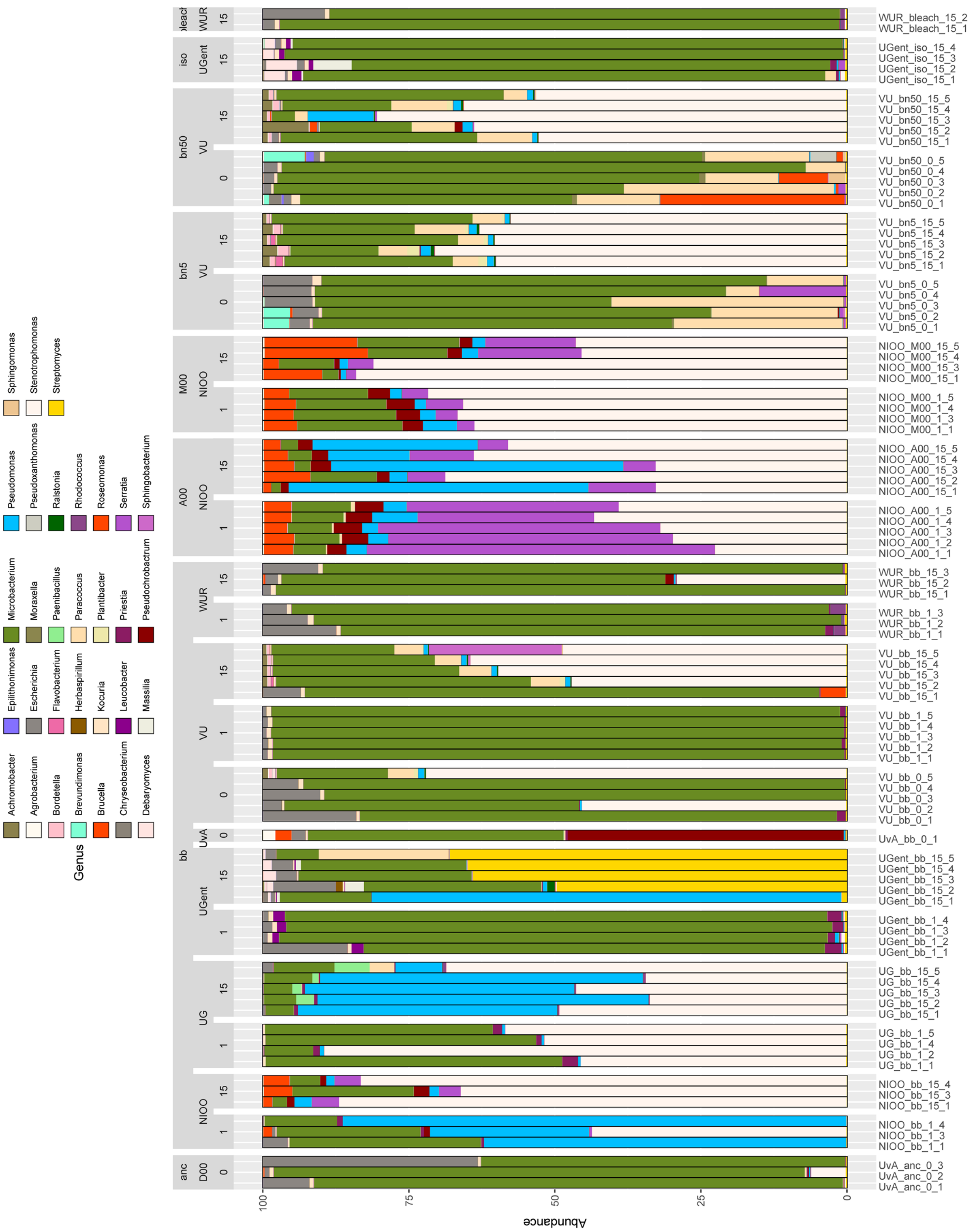


*__Figure S2: Institute-specific microbiome composition__. Relative abundance of bacterial genera across samples grouped by treatment (anc.: ancestral populations, bb: primary treatment, A00: androdioecious, M00: monoecious, bn5: bottleneck 5, bn50: bottleneck 50, iso: no mixing of lines, and bleach: initial bleaching step), institute (D00 ancestral population, NIOO, UG, UGent, VU, and WUR), and week (0, 1, and 15). The same information is provided in the label, and the last number indicates the specific replicate.*

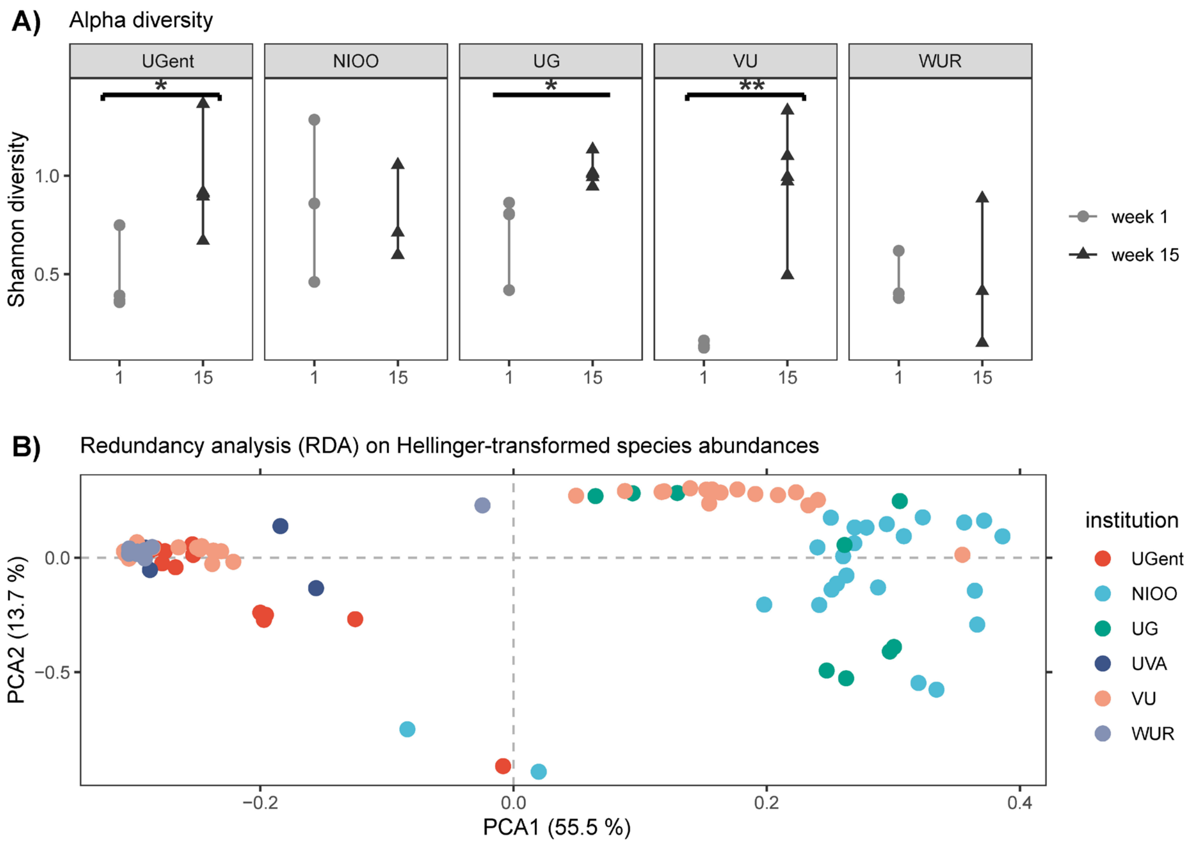


***Figure S3: Alpha and beta diversity of the microbiota.*** *A) The Shannon diversity is presented for each institute and week (week 1 in grey and week 15 in black). A significant increase in alpha diversity is found for UGent (p = 0.027), UG (p = 0.014), and VU (p = 0.009) (Table S3) B) A PCA plot based on the Hellinger-transformed species abundances showing the difference in microbiome composition between institutes.*

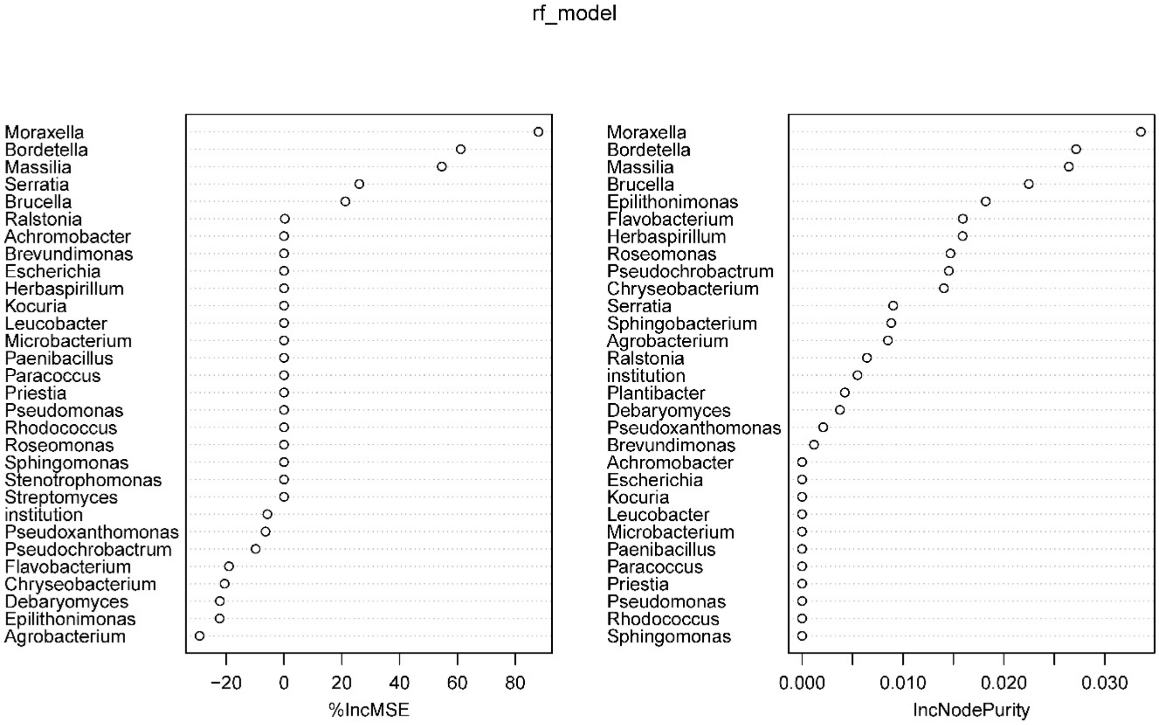


***Figure S4: Random Forest regression with institute and microbe data (always absent, lost, gained, always present) as explanatory variables.*** *Primary treatment samples with fitness change between week 1 and week 15 as response variable. %IncMSE is calculated as the increase in prediction error when randomly permuting a variable (high % increase in MSE suggests permuting that variable decreases model performance); IncNodePurity measures decrease in Residual Sum of Squares when 'splitting' on a variable, i.e. support for clustering when dividing the data by categories of the predictors (which institute or the presence/absence of bacterial taxa).*

TABLES

**Table S1. Overview of primary and secondary treatments conducted in the different institutes.**

| | **Primary (universal across institutes)** | **Secondary treatment level 1** | **Secondary treatment level 2** |
|---|---|---|---|
| **NIOO:** reproductive modes | **dioecious** (males and females) | **monoecious** (hermaphrodites) | **androdioecious** (hermaphrodites and males) |
| **UG** | | | |
| **UGent:** mixing of plates | **mixed** (three plates are mixed per replicate) | **isolated** (three plates are kept separately) | |
| **VU:** population bottlenecks | **no bottleneck** (500 nematodes to start initial population) | **moderate** (moderate bottleneck with 50 nematodes to start initial population) | **strong** (strong bottleneck with five nematodes to start initial population) |
| **WUR:** initial bleaching step | **no bleaching** | **bleaching** (an initial bleaching step at the start) | |

**Table S2: A) Chi-square statistics** with a significant institute effect and an interaction with time on the nematode growth rate. **B) Pairwise comparisons** adjusted for multiple comparisons using the Tukey method.

**A)** $\chi^2$ statistics: growth rate ~ week*institute + (1|replicate) + (1|measurement time)

| Independent variables | $X^2$ | df | *P* value |
|---|---|---|---|
| week | 0.4007 | 1 | 0.5267 |
| institute | 127.2001 | 4 | < 2.2e-16 |
| week:institute | 31.9760 | 4 | 1.935e-06 |

**B)** pairwise comparisons adjusted for multiple comparisons (Tukey method)

**contrast: week 15 - week 1**

| **institute** | estimate | SE | df | *Z* ratio | *P* value |
|---|---|---|---|---|---|
| NIOO | -0.023 | 0.035 | 187 | -0.659 | 0.5100 |
| UG | 0.105 | 0.029 | 187 | 3.609 | **0.0003** |
| UGent | -0.101 | 0.028 | 187 | -3.589 | **0.0003** |
| VU | 0.066 | 0.028 | 187 | 2.354 | **0.0186** |
| WUR | -0.030 | 0.044 | 187 | -0.694 | 0.4874 |

**within week 1**

| **contrast** | estimate | SE | df | *Z* ratio | *P* value |
|---|---|---|---|---|---|
| NIOO - UG | 0.215 | 0.044 | 187 | 4.931 | **<0.0001** |
| NIOO - UGent | 0.275 | 0.044 | 187 | 6.314 | **<0.0001** |
| NIOO - VU | 0.208 | 0.044 | 187 | 4.784 | **<0.0001** |
| NIOO - WUR | 0.297 | 0.050 | 187 | 5.981 | **<0.0001** |
| UG - UGent | 0.060 | 0.041 | 187 | 1.467 | 0.585 |
| UG - VU | -0.006 | 0.041 | 187 | -0.156 | 1.000 |
| UG - WUR | 0.082 | 0.047 | 187 | 1.725 | 0.418 |
| UGent - VU | -0.067 | 0.041 | 187 | -1.623 | 0.483 |
| UGent - WUR | 0.022 | 0.047 | 187 | 0.455 | 0.991 |
| VU - WUR | 0.088 | 0.047 | 187 | 1.860 | 0.339 |

**within week 15**

| **contrast** | estimate | SE | df | *Z* ratio | *P* value |
|---|---|---|---|---|---|

| | | | | | |
|---|---|---|---|---|---|
| NIOO - UG | 0.087 | 0.037 | 187 | 2.311 | 0.1414 |
| NIOO - UGent | 0.353 | 0.037 | 187 | 9.609 | **<0.0001** |
| NIOO - VU | 0.119 | 0.037 | 187 | 3.260 | **0.0098** |
| NIOO - WUR | 0.304 | 0.047 | 187 | 6.409 | **<0.0001** |
| UG - UGent | 0.267 | 0.033 | 187 | 8.184 | **<0.0001** |
| UG - VU | 0.033 | 0.033 | 187 | 1.009 | 0.8516 |
| UG - WUR | 0.217 | 0.044 | 187 | 4.904 | **<0.0001** |
| UGent - VU | -0.234 | 0.032 | 187 | -7.395 | **<0.0001** |
| UGent - WUR | -0.050 | 0.044 | 187 | -1.134 | 0.7886 |
| VU - WUR | 0.185 | 0.044 | 187 | 4.232 | **0.0002** |

***Table S3: Changes in time** for the different institutes for the **microbiota** data using the non-parametric Kruskall-Wallis rank sum test.*

| Institute | Contrast | Kruskal-Wallis $\chi^2$ | df | p-value |
|---|---|---|---|---|
| UGent | week 1 - week 15 | 4.860 | 1 | **0.027** |
| NIOO | week 1 - week 15 | 0.047 | 1 | 0.827 |
| UG | week 1 - week 15 | 6.000 | 1 | **0.014** |
| VU | week 1 - week 15 | 6.818 | 1 | **0.009** |
| WUR | week 1 - week 15 | 0.048 | 1 | 0.827 |

***Table S4: Pairwise comparisons** for the final fitness (population growth rates after 15 weeks) between different treatments within each institute. Results are adjusted for multiple comparisons using the Tukey method.*

| Institute | Contrast | Estimate | SE | df | t-ratio | p-value |
|---|---|---|---|---|---|---|
| UGent: effect of mixing | primary - no mixing | -0.276 | 0.033 | 47 | -8.438 | **<0.0001** |
| | primary - androdioecious | 0.032 | 0.300 | 47 | 0.108 | 0.9936 |
| | primary - monoecious | -0.890 | 0.322 | 47 | -2.764 | **0.0218** |
| NIOO: effect of reproductive mode | androdioecious - monoecious | -0.922 | 0.353 | 47 | -2.615 | **0.0315** |
| | primary - moderate | 0.465 | 0.253 | 83 | 1.843 | 0.1622 |
| VU: effect of population bottlenecks | primary - strong | 0.162 | 0.253 | 83 | -0.642 | 0.7973 |
| | moderate - strong | -0.303 | 0.281 | 83 | -1.079 | 0.5298 |
| WUR: effect of initial bleaching | primary - initial bleaching | -0.131 | 0.027 | 12 | -4.920 | **0.0004** |

***Table S5: The analysis of variance table of the best models for only the primary treatment.** The full model was the fitness (i.e. nematode growth rate) ~ week + institute + PC1 + PC2 + PC3, where the PCs are from the microbiota.*

| | Df | Sum Sq | Mean Sq | F-value | Pr(>F) |
|---|---|---|---|---|---|
| week | 1 | 0.04473 | 0.044734 | 0.9958 | **0.03153** |
| institute | 4 | 0.38404 | 0.096011 | 10.7223 | **7.151e-06** |
| PC3 | 1 | 0.04794 | 0.047940 | 5.3539 | **0.02633** |
| Residuals | 37 | 0.33131 | 0.008954 | | |

***Table S6: The analysis of variance table of the best models for the primary and secondary treatments.*** *The full model was the fitness (i.e. nematode growth rate) ~ week + institute + treatment + PC1 + PC2 + PC3, where the PCs are from the microbiota.*

| | Df | Sum Sq | Mean Sq | F-value | Pr(>F) |
|---|---|---|---|---|---|
| week | 1 | 0.45464 | 0.45464 | 39.8987 | 2.892e-08 |
| institute | 5 | 0.70137 | 0.14027 | 12.3102 | 2.241e-08 |
| treatment | 6 | 0.27282 | 0.04547 | 3.9904 | 0.001872 |
| PC1 | 1 | 0.05929 | 0.05929 | 5.2031 | 0.025884 |
| PC3 | 1 | 0.06338 | 0.06338 | 5.5622 | 0.021418 |
| Residuals | 64 | 0.72928 | 0.01140 | | |
