## Appendix I for "Additive effects of environmental and demographic variation shape the repeatability of evolution across replicated experiments"

Title: Additive effects of environmental and demographic variation shape the repeatability of evolution across replicated experiments.

### 1. Preparation of media and buffers

Media and buffers are adapted from Stiernagle (2006).

##### *a. Preparation of the Nematode Growth Medium (NGM)*

Ingredients:

- 3 g NaCl
- 17 g Agar
- 2.5 g Peptone
- 1 mL cholesterol (5 mg/mL in ethanol; do not autoclave; store at -20°C)
- 25 mL 1M $KPO_4$ buffer pH 6.0 (108.3 g $KH_2PO_4$, 35.6 g $K_2HPO_4$, $H_2O$ to 1 litre; bring to pH 6.0; autoclave for 15 minutes at 120°C; store at room temperature)
- 1 mL 1M $MgSO_4$ (12 g $MgSO_4$ and $H_2O$ to 100 mL; autoclave 15 minutes at 120°C; store at room temperature)
- 1 mL 1M $CaCl_2$ (14.7g $CaCl_2.2H_2O$ and $H_2O$ to 100 mL; autoclave 15 minutes at 120°C; store at room temperature)

Preparation:

- Mix 3 g NaCl, 17 g agar, and 2.5 g peptone in a 1 litre bottle
- Add 975 mL $H_2O$
- Autoclave (15 min, 120°C)
- Cool flask to 55°C in water bath or stove
- Add 1 mL 1 M $CaCl_2$, 1 mL 5 mg/mL cholesterol in ethanol, 1 mL 1 M $MgSO_4$ and 25 mL 1 M $KPO_4$ buffer. Swirl to mix well.

##### *b. Preparation of LB agar*

- 10 g Bacto-tryptone
- 5 g Bacto-yeast
- 5 g NaCl
- 15 g agar
- $H_2O$ to 1 litre
- Bring to pH 7.5
- Autoclave (15 minutes at 120°C)

#### 2. Preparation of bacterial stocks

Preparation of L broth

- 10 g Bacto-tryptone
- 5 g Bacto-yeast
- 5 g NaCl
- $H_2O$ to 1 litre
- Bring to pH 7.0 using 1M NaOH
- Autoclave (15 minutes at 120°C)
- Store at room temperature

Protocol

- Pick a colony from the stock plate using a sterile tip
- Drop the tip in 10 mL of L broth
- Grow for 20 hours at 37°C while shaking at 400 rpm
- Store the cultures in the fridge for maximum 4 weeks

Title: Additive effects of environmental and demographic variation shape the repeatability of evolution across replicated experiments.

### 3. Washing nematodes off plates

Preparation of buffer

*M9*

- 5.8 g $Na_2HPO_4.7H_2O$
- 3.0 g $KH_2PO_4$
- 5.0 g NaCl
- $H_2O$ to 1 litre
- Autoclave (15 minutes at 120°C)
- Add 1ml sterile 1M $MgSO_4.7H_2O$
- Store at room temperature

*S-buffer*

- 0.05 M $K_2HPO_4$
- 0.05 M $KH_2PO_4$
- 5.85 g NaCl
- $H_2O$ to 1 L
- Autoclave (15 minutes at 120°C)
- Store at room temperature

Protocol

- Pipette 2mL buffer onto plate
- Swirl and pipet up and down a few times
- Transfer to Eppendorf tube
- Let nematodes sink to bottom for 15 minutes in fridge OR centrifuge 10 seconds at 6000 rpm (20°C)
- Remove supernatant

Side notes: not all nematodes will be removed as the nematodes that have crawled into the NGM will remain (leaving the buffer for a while on the plate will encourage nematodes to leave the NGM).

Title: Additive effects of environmental and demographic variation shape the repeatability of evolution across replicated experiments.

### 4. Freezing of nematodes

#### *a. Soft agar freezing*

Preparation soft agar

- 0.58 g NaCl
- 0.68 g $KH_2PO4$
- 30 g glycerol
- 560 µL 1M NaOH
- 0.4 g agar
- $H_2O$ to 100 mL
- Autoclave (15 minutes at 120°C)
- Melt agar before use in microwave and keep warm in water bath (50°C)

Protocol

- Wash the nematodes off the plate following the protocol (Appendix III)
- Transfer suspension to 2mL cryo-vial
- Put on ice for 20 minutes
- Remove supernatant partially (leave 0.5 mL buffer in the cryo-vial)
- Add 0.5 mL soft agar freezing solution and shake
- Place cryo-vial in Mr. Frosty or Styrofoam box with cotton wool
- Freeze in -80°C for 24h
- Transfer cryo-vial to normal -80°C cryo-box storage

#### *b. Liquid agar freezing*

Preparation liquid freezing solution

- 5.8 g NaCl
- 50 mL 1M $KH_2PO_4$ (pH 6.0)
- 240 mL glycerol (15% glycerol concentration in final solution)
- 710 mL $H_2O$
- Autoclave (15 minutes at 120°C)
- Add 30 µL sterile 1M $MgSO_4$ per 100 mL liquid freezing solution

Protocol

- Wash the nematodes off the plate following the protocol (Appendix III)
- Transfer suspension to 2mL cryo-vial
- Leave cryo-vial at room temperature for 20 minutes to let nematodes sink to bottom

- Remove supernatant partially (leave 0.5 mL buffer in the cryo-vial)
- Add 0.5 mL liquid freezing solution and mix by inverting tube
- Place cryo-vial in Mr. Frosty or Styrofoam box with cotton wool
- Freeze in -80°C for 24h
- Transfer cryo-vial to normal -80°C cryo-box storage

Title: Additive effects of environmental and demographic variation shape the repeatability of evolution across replicated experiments.

#### 5. Differences between institutes

**Table A1: Details on deviations in the experimental procedure across institutes.** The columns indicate the different institutes (NIOO: Netherlands Institute of Ecology NIOO-KNAW, UG: University of Groningen, UGent: Ghent University, VU: Vrije Universiteit Amsterdam, WUR: Wageningen University & Research).

| | **NIOO** | **UG** | **UGent** | **VU** | **WUR** |
|---|---|---|---|---|---|
| **Institute set-up** | | | | | |
| **Did you work in a flow cabinet or under a flame?** | | | | | |
| **Flow cabinet** | X | X* | X | X | X |
| **Flame** | | X* | X | X | |
| *Transfers were done next to the flame for the first three weeks, afterwards in a flow cabinet. | | | | | |
| **Were the nematodes kept in a climate cabinet or room?** | | | | | |
| **Climate cabinet** | X | | X* | | X |
| **Climate room** | | X | X* | X | |
| *The climate cabinets were in a room with temperature regulation. | | | | | |
| **Was there light in the climate cabinet or room?** | | | | | |
| **Yes (L:D)** | 16:08 | | | 16:08 | |
| **No** | | X | X | | X |
| **What was the relative humidity in the climate cabinet or room?** | | | | | |
| **% humidity** | | 70% | | 75% | |
| **Not measured** | X | | X* | | X |
| *A box with water was placed inside the climate cabinet for humidity. | | | | | |
| **Bacterial lawns** | | | | | |
| **When did you prepare new *P. megaterium* cultures?** | | | | | |
| **How often?** | Every 4 weeks | Every 4 weeks | Every 2 weeks* | Every 3 weeks | Every 4 weeks |
| *New *P. megaterium* cultures were the first time made after one month, afterwards every two weeks. | | | | | |
| **How many days before transfer did you seed the NGM plates with *P. megaterium*? OR was this done once for a few transfers ('once')?** | | | | | |
| **Days before** | 1-2*1 | 1 | | | 0 |
| **Once** | | | monthly | monthly*3 | |
| *1Last transfer the plates were seeded a week in advance.<br>*2Initially plates were seeded two days before transfer, afterwards on the same day.<br>*3Once per month for the first 9 weeks, from the 10th week onward every two weeks. | | | | | |
| **How did you grow the bacteria on the plates? (RT: room temperature)** | | | | | |
| **1 hour RT** | | | | | X |
| **1 day RT** | X | | | X | |
| **2 days RT** | | | | | |
| **Overnight in a stove (37°C)** | | X | X | | |
| **Where did you store the NGM plates without bacteria? (RT: room temperature)** | | | | | |
| **Fridge (4°C)** | X | | X | X | X |
| **Other** | | X (16°C) | | | |
| **Washing the nematodes off the plates** | | | | | |

Title: Additive effects of environmental and demographic variation shape the repeatability of evolution across replicated experiments.

| | | | | | |
|---|---|---|---|---|---|
| **Which buffer was used?** | | | | | |
| **S-buffer** | X | | X | X | |
| **M9** | | X | | | X |
| **How many mL buffer did you use?** | | | | | |
| **Volume (mL)** | 2 mL | 2 mL | 3 mL[*] | 3 mL | 2 mL |
| [*]In the side experiments between 2 and 2.5 mL | | | | | |
| **How long was the buffer on the plate?** | | | | | |
| **< 1 min.** | X | | | | X |
| **1 – 3 min.** | | X | X | X | |
| **How many times did you swirl the plate?** | | | | | |
| **1 – 3** | X | X | | | X |
| **4 – 6** | | | X | X | |
| **How many times did you pipet up and down?** | | | | | |
| **1 – 2** | | | X | | |
| **3 – 5** | X | | | | X |
| **> 5** | | X | | X | |
| **Centrifuge or let the nematodes sink to bottom in a fridge during washing?** | | | | | |
| **Centrifuge** | | X | | | |
| **Sink in fridge** | | | X | X | |
| **Sink at RT** | X | | | | X |
| **Counting nematodes** | | | | | |
| **What volume did you count?** | | | | | |
| **Volume (µL)** | 5 µL | 1-2 µL | 5 µL | 2 µL | 2 µL |
| **How many drops did you count?** | | | | | |
| **Number of drops** | 5 | 3 | 1[*] | 3 | 3 |
| [*]In case the count was very low, a second droplet was counted to verify. | | | | | |
| **Did you dilute before counting?** | | | | | |
| **Yes** | X | X | X | | |
| **No** | | | | X | X |
| **Freezing** | | | | | |
| **What proportion of the nematodes did you freeze (after transfer to fresh plates)?** | | | | | |
| **< 25%** | | | | | |
| **25 – 50%** | | | X | | |
| **100%** | X | X | | X | X |
| **Over how many Eppendorf or cryo-tubes were the nematodes divided?** | | | | | |
| **Number of tubes** | 3 | 5 | 2[*2] | 1 | 1 |
| [*1]The nematodes from the different replicas have been combined in the same tube; plan to redo the experiment<br>[*2]At weeks 5, 10, and 15 extra samples were frozen | | | | | |
| **Did you use Mr. Frosty, a Styrofoam box or nothing to freeze the nematodes?** | | | | | |
| **Mr. Frosty** | | | X | X | |
| **Styrofoam box** | X | | | | |
| **Nothing** | | X | | | X |
| **Liquid or soft agar freezing** | | | | | |
| **Liquid freezing** | | X | | | |
| **Soft agar freezing** | X | X | X | X | X |
| **Other** | | | | | |
| **Are there ‘population’ counts available? Note: these counts were to transfer nematodes and not scaled towards the total volume of suspension.** | | | | | |

| | | | | | |
|---|---|---|---|---|---|
| **Yes** | X | | | X | X |
| **No** | | X | X* | | |
| *Only from the last week | | | | | |
| **Was it necessary to restart from a previous week?** | | | | | |
| **Yes (which week)** | | | | | |
| **No** | X | X | X | X | X |

Title: Additive effects of environmental and demographic variation shape the repeatability of evolution across replicated experiments.

### 6. Validation of the data

#### *a. Correlation between manual and BioSorter counts.*

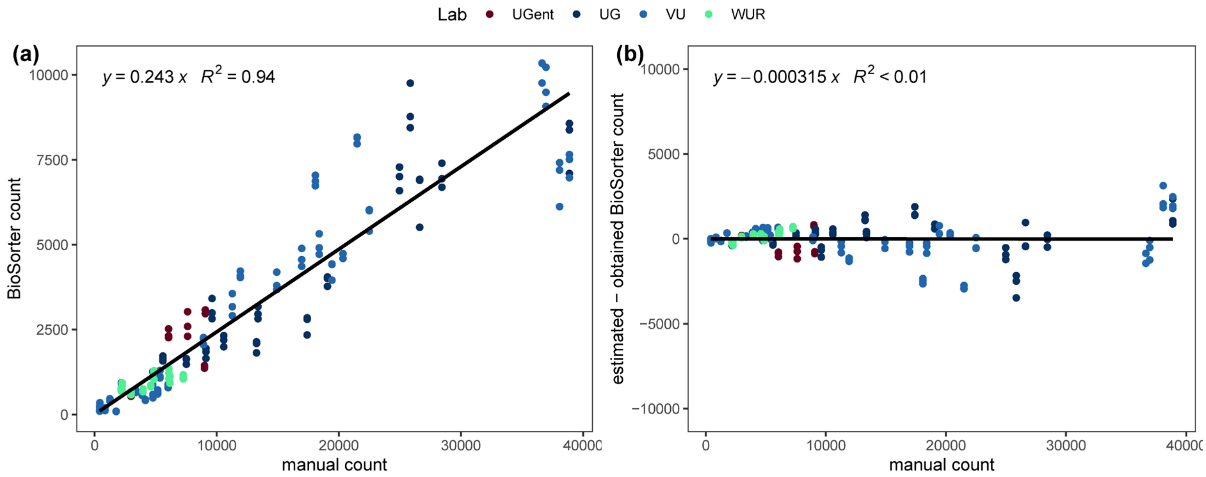

**Figure A1: Correlation between manual and BioSorter counts.** (a) The x-axis shows the manual counts, while the y-axis gives the results from the flow cytometer or BioSorter. Each dot represents the same technical replicate for which the population size was assessed. The linear regression through the origin is indicated with the solid line and the equation is presented in the upper left corner. (b) The manual counts are given on the x-axis, while the y-axis represents the difference between the estimated counts based on the equation in (a) and the obtained counts from the flow cytometer.

Title: Additive effects of environmental and demographic variation shape the repeatability of evolution across replicated experiments.

*b. Growth curves as validation for measurement of population sizes at day 7.*

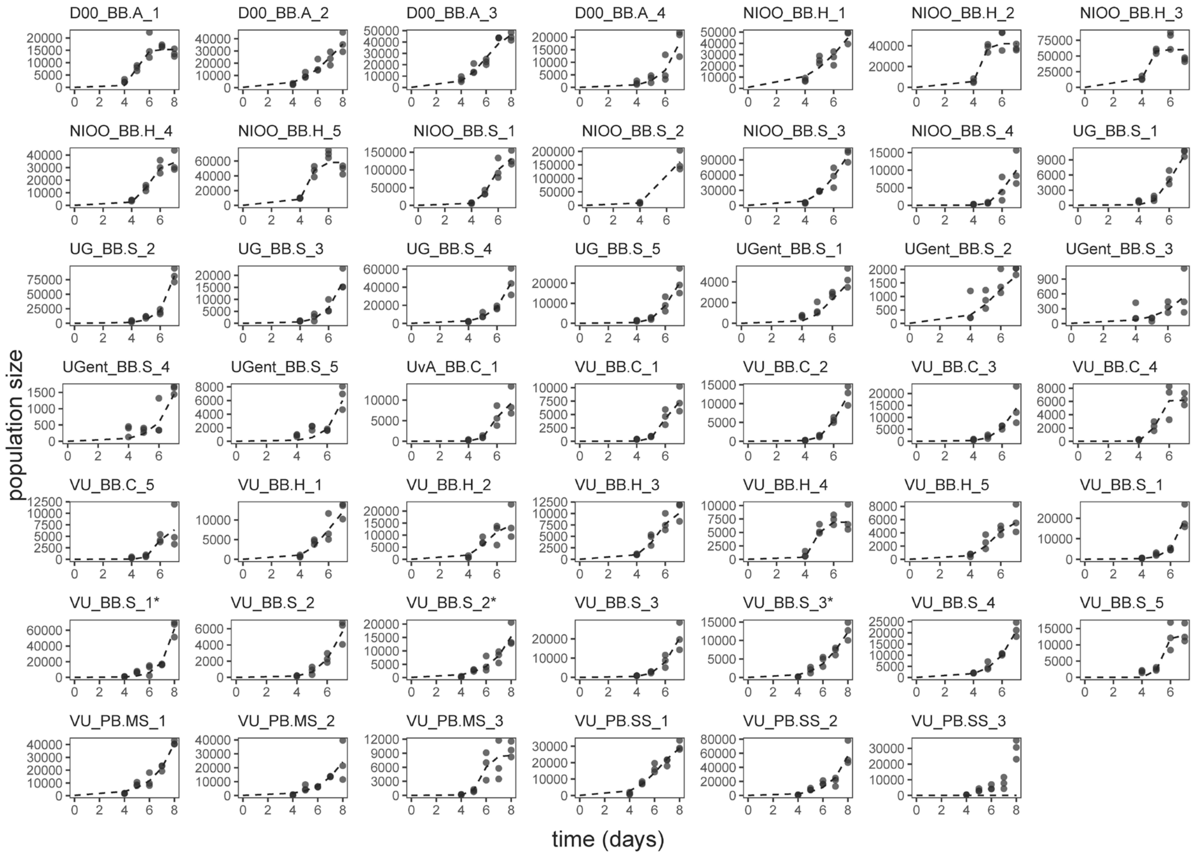

**Figure A2: growth curves as validation for the measurement of population sizes at day 7.** Each panel represents a replicate for which population size measurements (y-axis) were assessed for 7 or 8 days (x-axis). Per day three technical replicates were measured. In the manuscript we included data from the population sizes (the number of nematodes after one week of growth on the novel host plant) as proxy of fitness as this is the closest to their real fitness due to our experimental setup (the plates were refreshed every week). To further validate this proxy we also calculated the growth rate for some populations to see whether the growth is not decreasing again at day seven, but still growing or stabilizing.

Title: Additive effects of environmental and demographic variation shape the repeatability of evolution across replicated experiments.

*c. Population sizes partitioned across measurement days.*

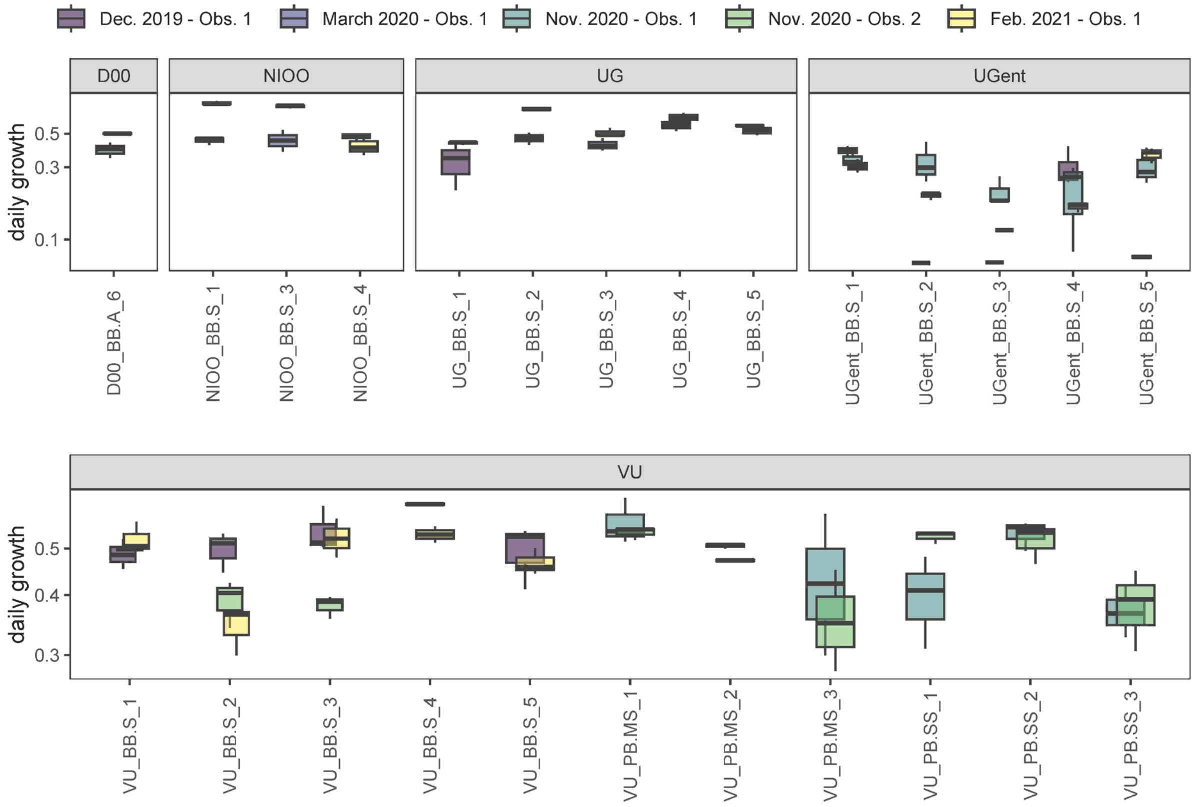

**Figure A3: Population size after 7 days of growth on *P. megaterium* before and after selection, partitioned across measurement days.** Each box-plot shows the distribution of three measurements taken at a given day (colours) by either observer 1 (purple, dark blue, blue, yellow) or observer 2 (green) and each facet shows the different institutes (D00: ancestor, NIOO: NIOO-KNAW, the Netherlands Institute of Ecology, UG: University of Groningen, UGent: Ghent University, VU: Vrij Universiteit Amsterdam). Labels: BB.A: ancestral population, BB.S = primary treatment at week 15, PB.MS = moderate population bottleneck at week 15, PB.SS = strong population bottleneck at week 15.
